# Increased wheat protein content via introgression of a HMW glutenin selectively reshapes the grain proteome

**DOI:** 10.1101/2020.10.02.324137

**Authors:** Hui Cao, Owen Duncan, Shahidul Islam, Jingjuan Zhang, Wujun Ma, A. Harvey Millar

## Abstract

Introgression of a high molecular weight glutenin subunit (HMW-GS) gene, *1Ay21\**, into commercial wheat cultivars increased overall grain protein content and bread-making quality by unknown mechanisms. As well as increased abundance of 1Ay HMW-GS, 115 differentially expressed proteins (DEPs) were discovered between three cultivars and corresponding introgressed near-isogenic lines (NILs). Functional category analysis showed that the DEPs were predominantly other storage proteins, and proteins involved in protein synthesis, protein folding, protein degradation, stress response and grain development. Nearly half the genes encoding the DEPs showed strong co-expression patterns during grain development. Promoters of these genes are enriched in elements associated with transcription initiation and light response, indicating a potential connection between these *cis*-elements and grain protein accumulation. A model of how this HMW-GS enhances the abundance of machinery for protein synthesis and maturation during grain filling is proposed. This analysis not only provides insights into how introgression of the *1Ay21\** improves grain protein content, but also directs selection of protein candidates for future wheat quality breeding programmes.

**One sentence summary:** Introgression of the 1Ay21* HMW-GS increases wheat grain protein content and improves bread-making quality in association with a broad reshaping of the grain proteome network.

## INTRODUCTION

Hexaploid wheat (*Triticum aestivum* L. 2n = 6x = 42, AABBDD) is cultivated worldwide due to its value as a staple food and protein source. Apart from the relatively high starch content (60-70% of the whole grain dry weight), wheat grain protein content varies from 8-15%, and this determines bread-making qualities (Shewry, 2009). Grain protein content largely consists of storage proteins including glutenins and gliadins, of which glutenins determine the dough viscoelasticity and elasticity while gliadins contribute to dough extensibility (Shewry and Halford, 2002; Rasheed et al., 2014). Glutenins can be further divided into two groups, namely high molecular weight glutenin subunits (HMW-GS, 70-140 kDa) and low molecular weight glutenin subunits (LMW-GS, 30-80 kDa) (Shewry et al., 1992; Shewry et al., 2002).

Generally, HMW-GS encoding genes are located at the *Glu-1* loci on the long arms of chromosomes 1A, 1B and 1D. Each *Glu-1* locus consists of two tightly linked genes with one gene encoding an x-type small subunit and the other gene encoding an y-type large subunit, named according to molecular weight differences between the encoded HMW-GS proteins (Payne, 1983; Nakamura, 1999). Among these six subunits, three subunits (Glu-1Bx, Glu-1Dx and Glu-1Dy) are always expressed in bread wheat cultivars while two subunits (Glu-1Ax and Glu-1By) are sometimes expressed, but the Glu-1Ay subunit is rarely if ever expressed (Waines and Payne, 1987). As a result, most common bread wheat cultivars usually have three to five expressed HMW-GSs. However, the *Glu-1Ay* gene has been found to be functionally expressed in wild diploid and tetraploid wheats (Ciaffi et al., 1993; Jiang et al., 2009).

Grain protein content and composition are important traits in global bread wheat breeding programmes due to the continuous growth in demand for specified industrial end-use (Shewry, 2007; Kiszonas and Morris, 2018). Glutenin in particular, has become one of the main breeding targets because of its critical role in determining bread-making quality. For example, it has been previously reported that allelic variation of HMW-GS contributes to about 45-70% of the variation in bread-making quality, although it only contributes to about 10-12% of total grain protein content (Branlard and Dardevet, 1985; Payne et al., 1987; Liu et al., 2007). Furthermore, y-type HMW-GSs are generally considered to be more valuable for dough quality improvement than x-type HMW-GSs because they are longer and possess more cysteine residues for forming inter- and intra-molecular disulphide bonds during dough development (Shewry and Halford, 2002). Genetic engineering strategies have been widely used to improve protein content and composition in wheat breeding, which includes the use of conventional cross breeding processes to integrate additional genes into background genotypes to produce extra protein functionality and increase protein content (Shewry, 2007). Margiotta et al. (1996) reported the identification of active *1Ax* and *1Ay* genes in several Swedish bread wheat lines, and a year later Rogers et al. (1997) demonstrated that 1Ay HMW-GS had positive effects on bread-making quality, such as improved dough stability during mixing and enhanced gluten strength (Margiotta et al., 1996; Rogers et al., 1997). Since then, attention has been drawn to understanding the biological function of 1Ay HMW-GS and to estimate its value in wheat breeding programmes. Li et al. (2006) reported the investigation of HMW-GS variation amongst 205 cultivated emmer accessions, which led to the successful identification of novel 1Ay subunits and the recommendation of emmer accessions as a valuable genetic resource for quality improvement of common wheat (Li et al., 2006). Another study, focusing on genetic variation of HMW-GSs in 1051 accessions from 13 *Triticum* subspecies including diploid, tetraploid and hexaploid wheat, identified the expression of 1Ay subunits in *T*.*urartu* and suggested these unique HMW-GS alleles could be further utilized through direct hybrid production for quality improvement of common wheat (Xu et al., 2009).

Recently, Roy and co-workers integrated an expressed 1Ay allele from an Italian line into Australian commercial cultivars using a conventional breeding approach, and observed better dough rheological properties and bread-making quality without significant changes in agronomic traits. Improved quality traits observed including higher protein content, Glu/Gli ratio and UPP%, stronger dough strength and better water absorption (Roy et al., 2018; Roy et al., 2020). It is well known that improving grain yield and grain protein content simultaneously is difficult due to their inverse correlation (Iqbal et al., 2007). However, the introgression of the 1Ay21* HMW-GS in the Lincoln cultivar background, resulted in an increase in both grain yield and protein content in glasshouse and field conditions (Roy et al., 2020). Yu et al cloned and characterized the *Glu-1Ay* gene sequence identified in the Italian line and suggested that it has a close evolutionary relationship with Ay genes of *T. urartu* (Yu et al., 2019).

However, how the introgressed *Glu-1Ay* gene increases bread wheat protein content without changing other agronomic traits is still unknown. Here we have undertaken a systematic proteome analysis for three Ay introgressed near isogenic lines (NIL) in Australian cultivar backgrounds. Our findings provide new insights into which grain proteins are increased or decreased in abundance in introgressed lines and proposes a common link between them.

## RESULTS

### Alteration in HMW-GSs composition and total grain protein content

Through a conventional cross breeding process, an expressed *Glu-Ay* allele from an Italian wheat line was successfully introgressed into three Australian bread wheat cultivars, namely Gregory, Bonnie Rock and Yitpi. In each cross *Ax1* or *Ax1*\* and *Ay*^*NE*^ (Not Expressed) genes were replaced by *Ax21* and *Ay21\** genes, respectively (Supplemental Table S1A) (Roy et al., 2018). From these crosses, near isogenic lines (NILs) were developed in each cultivar. In this study, three lines including one parental cultivar and two Ay integrated NILs for each of the three cultivars were subjected to a quantitative proteome analysis. Total protein extracts from mature grain samples were extracted, digested with trypsin and analysed by LC-MS/MS. This yielded large spectral libraries of MS/MS data and comparable datasets of precursor ion intensities that were amenable to label free quantification.

Initially we sought to use these data to confirm the presence of the Ay HMW-GS protein in NILs and measure the abundance of other HMW-GSs in parental cultivars by searching the data against the peptide sequences listed in Supplemental Table S1A. These data were expressed as iBAQ (Intensity Based Absolute Quantification) values which allow comparison between the same protein in different genotypes and also between different protein types within the genotype (Tyanova et al., 2016). This confirmed that five HMW-GSs were detected in parental cultivars and peptides matching to the sixth Ay HMW-GS protein were only detected in significant abundance in NILs (Fig. 1). The Ay HMW-GS amount detected in NILs was approximately 15% of all y-type HMW-GS protein abundance and approximately 6% of total HMW-GS content. There was some background detection of Ay HMW-GS like peptides in Gregory, but these were only approximately 11% of the peptide abundances of Ay HMW-GS in the Gregory NILs (Supplemental Table S1). Apart from the presence of Ay HMW-GS, no significant absolute abundance changes were detected for peptides matching the other five HMW-GSs in any of the NILs compared to their corresponding parental cultivar. This evidence that introgression of Ay HMW glutenin subunit did not affect the absolute abundance of the other five HMW-GSs is consist with previous results based on HPLC analysis of HMW-GS subunits (Roy et al., 2018).

**Figure 1.**
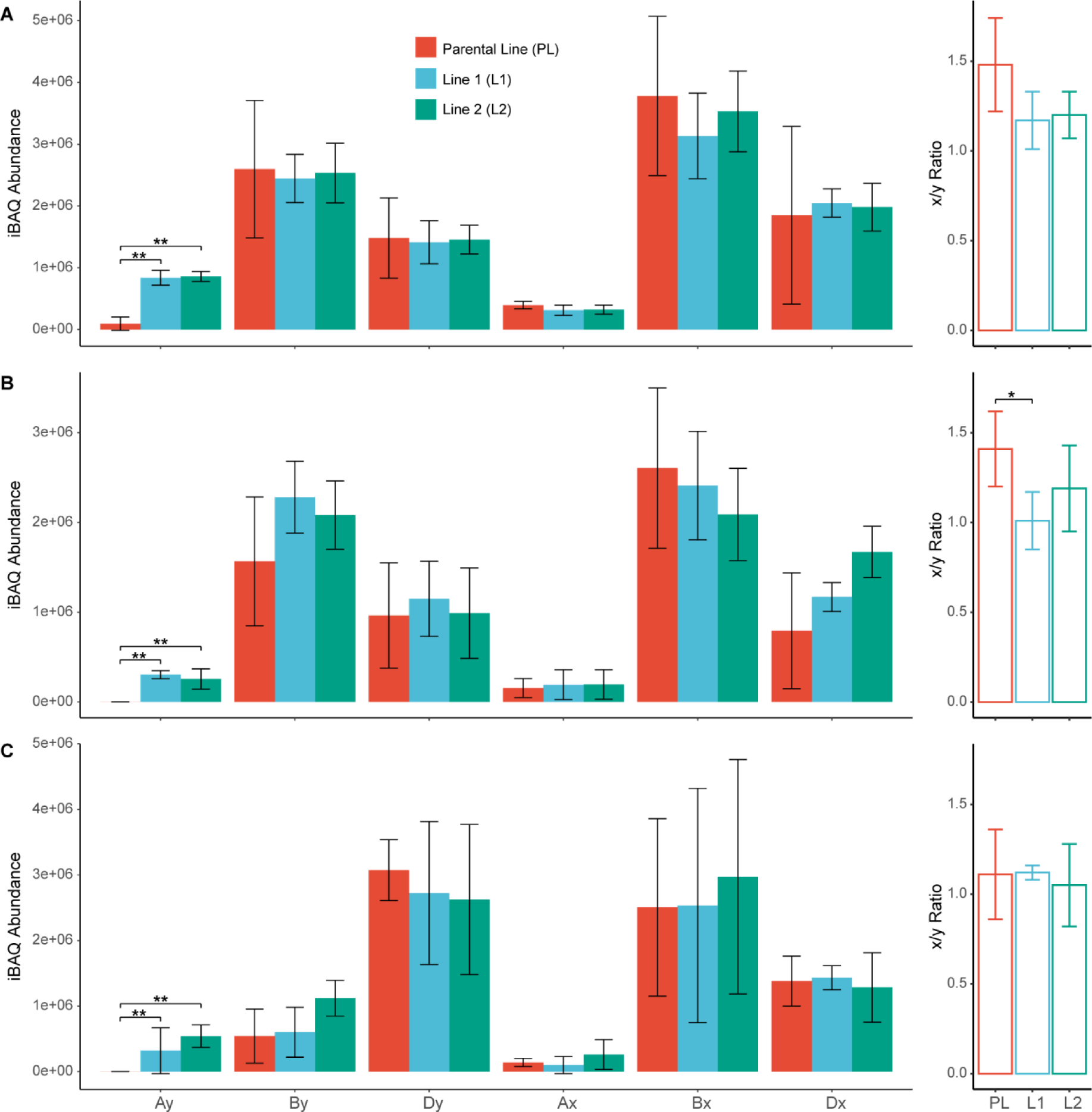
HMW-GSs abundance in parental and Ay integrated lines. HMW-GSs abundance estimated through Intensity Based Absolute Quantification (iBAQ) analysis (MaxQuant) for four biological replicates. Detailed results are shown in Supplemental Table S1. The hollow bars on the right panel represent the ratio of x-type and y-type HMW-GSs for each line. A, HMW-GSs absolute abundance for cv. Gregory lines. B, HMW-GSs absolute abundance for cv. Bonnie Rock lines. C, HMW-GSs absolute abundance for cv. Yitpi lines. Significant abundance changes between lines are marked by asterisks, * ⩽ 0.05; ** ⩽ 0.01.

The x-type to y-type HMW-GS ratio is critical to dough quality and bread-making quality with lower numbers correlating with higher quality (Leon et al., 2010; Roy et al., 2018). Our label free quantitative analysis showed x-type/y-type ratio of 1.48, 1.41 and 1.11 for Gregory, Bonnie Rock and Yitpi parental cultivars, respectively (Fig. 1). This ratio marginally decreased in the presence of Ay HMW-GS in NILs, although only one was statistically significant, the decrease found in Bonnie Rock integrated line 1 (Fig. 1B). We also found that some of the glutenin subunits, including Ay HMW-GS, Bx and Dx HMW-GSs in Bonnie Rock integrated line2 and Dy HMW-GS in Yitpi integrated line2, significantly decreased in their relative contribution to the whole HMW-GS pool (Supplemental Table S1B-D). Overall, this showed that while the introgression of Ay HMW-GS significantly changed the composition of the HMW-GSs pool in some lines, the absolute abundance of all HMW-GSs and x-type to y-type HMW-GS ratio of majority lines did not statistically change.

It has been reported that the introgression of Ay HMW-GS can increase the total protein content and gluten content of the grain (Roy et al., 2018; Roy et al., 2020). Total protein content on a dry weight basis showed a significant increase in grain protein in 5 of the 6 integrated NIL lines compared to their parental cultivar with the amount of grain protein on a grain weight basis being approximately 1% (w/w) higher in the NIL lines (Table 1). Total gluten content on a protein basis increased by about 5% in the NILs. Given total HMW glutenin content contributes 2-3% to grain weight (Zorb et al., 2018), the increased amount of Ay HMW-GS alone would contributed to less than 0.2% (w/w) of the total protein amount increase observed, suggesting that a substantial proportion of the observed increase was not derived from the extra Ay HMW-GS but from other grain proteins increasing in abundance.

**Table 1.** Changes in total wheat grain protein and gluten content in Ay integrated lines. The averaged % protein (w/w) on a grain weight basis and % gluten content (w/w) on a total protein basis with standard deviation (± sd) were presented (*n = 3*). Significant changes were determined using t-test *p ⩽ 0.05, **p ⩽ 0.01.

| Cultivar | Line | Total Protein Content |  | Gluten Content |  |
| --- | --- | --- | --- | --- | --- |
|  |  | Protein (%) | Net Increase (%) | Gluten (%) | Net Increase (%) |
| Gregory | Parent | 11.76 ± 0.36 | -- | 22.07 ± 0.87 | -- |
| | Line 1 | 12.40 ± 0.10 | 0.64 * | 22.80 ± 0.91 | 0.73 |
| | Line 2 | 12.06 ± 0.23 | 0.30 | 23.93 ± 0.31 | 1.86 * |
| Bonnie Rock | Parent | 11.77 ± 0.29 | -- | 21.97 ± 0.42 | -- |
| | Line 1 | 12.47 ± 0.15 | 0.70 * | 27.30 ± 0.56 | 5.33 ** |
| | Line 2 | 12.33 ± 0.17 | 0.56 * | 27.42 ± 0.26 | 5.45 ** |
| Yitpi | Parent | 12.10 ± 0.31 | -- | 25.07 ± 0.62 | -- |
| | Line 1 | 13.43 ± 0.28 | 1.33 ** | 30.39 ± 0.64 | 5.32 ** |
| | Line 2 | 13.23 ± 0.11 | 1.13 ** | 30.03 ± 0.80 | 4.96 ** |

### Impacts of the introgression of 1Ay21* HMW glutenin subunit on grain proteome profiles

To find out which other wheat grain proteins changed in abundance and were contributing to the protein content increase (Table 1), we conducted a systematic analysis of the proteomics data from both parental and integrated lines for all three cultivars. There were high correlations between peptide relative abundances measured across the four bio-replicates. The Pearson coefficient correlation ranged from 0.70 to 0.90 between replicates, with an average value of 0.79. In total, over 11,000 quantifiable peptides were detected across three cultivars and these mapped to 683, 648 and 722 proteins from EGA Gregory, Bonnie Rock and Yitpi, respectively (Supplemental Table S2). A principal component analysis of protein relative abundance showed that the three cultivars could be separated by a single principal component explaining nearly 62% of the variation, while there were only minor variations observed between NILs of the same cultivar (Fig. 2A). PCA analysis of data from each individual cultivar indicated that 50%-60% of the variation could be explained by two components and the parental cultivars were clearly separated from the integrated lines (Fig. 2B). Further hierarchical clustering analysis of protein relative abundance data showed that the Ay integrated lines were more similar to each other than to the parental cultivar, for both Bonnie Rock and Gregory. Similarly, replicates of the same line had a closer relationship to each other than to other lines with some exceptions found in Gregory samples (Fig. 2C).

**Figure 2.**
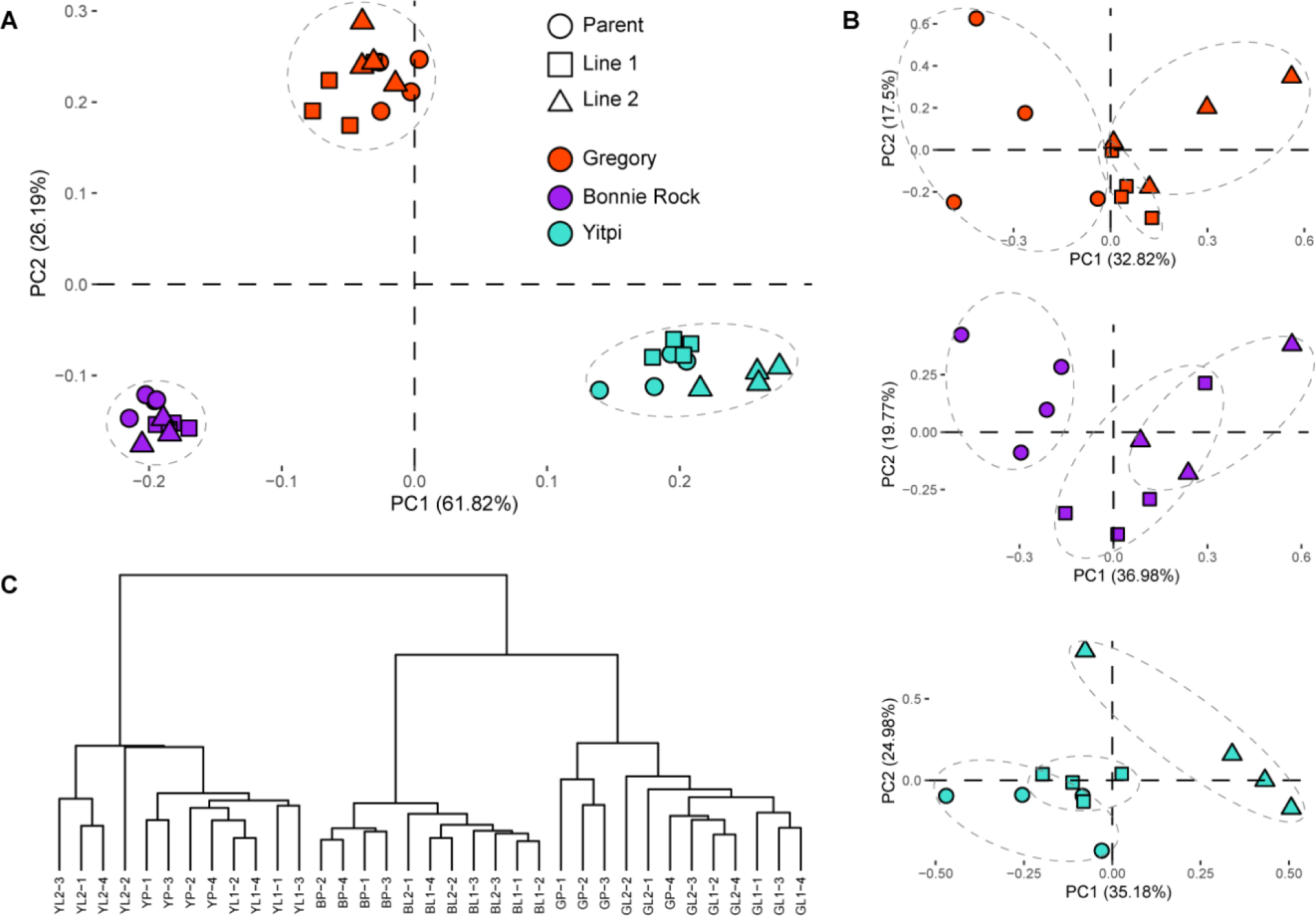
Principal component analysis of protein abundances from parental and Ay integrated lines. A, the PCA for all 35 samples from three cultivars. B, the individual PCA for each cultivar. C, the dendrogram of hierarchical clustering analysis for the same data used in A. A data set of 185 proteins that have relative protein abundance value across all 35 samples was used in A, while the data set in B included 353 proteins for cv. Gregory, 404 for cv. Bonnie Rock and 432 for cv. Yitpi. The full data used in this analysis is provided in Supplemental Table S2D-G. The dashed circles outline biological replicates for each cultivar in A and each line in B. YP and YL represent the cv. Yitpi parental and integrated line, while G and B represent cv. Gregory and cv. Bonnie Rock, respectively.

Focusing on proteins found in a parental cultivar and that had the same pattern of change in abundance across both of the integrated NILs, we identified 34 DEPs that increased in abundance and 13 DEPs that decreased in abundance in Gregory (Fig. 3A). For Bonnie Rock and Yitpi these numbers were 14 and 16, and 6 and 10, respectively (Fig. 3B and C). While ribosome protein subunits where found as DEPs both increasing and decreasing in abundance, storage proteins typically increased in abundance while biotic and abiotic stress response proteins typically decreased in abundance. This approach to analysis, however, filtered out present/absent differences that could only be uniquely detected in either parental cultivar or the NILs. Including such present/absent proteins (Fig. 3D) more than doubled the number of DEPs that could be considered in the analysis (Supplemental Fig. S2 & Supplemental Table S3H and I).

**Figure 3.**
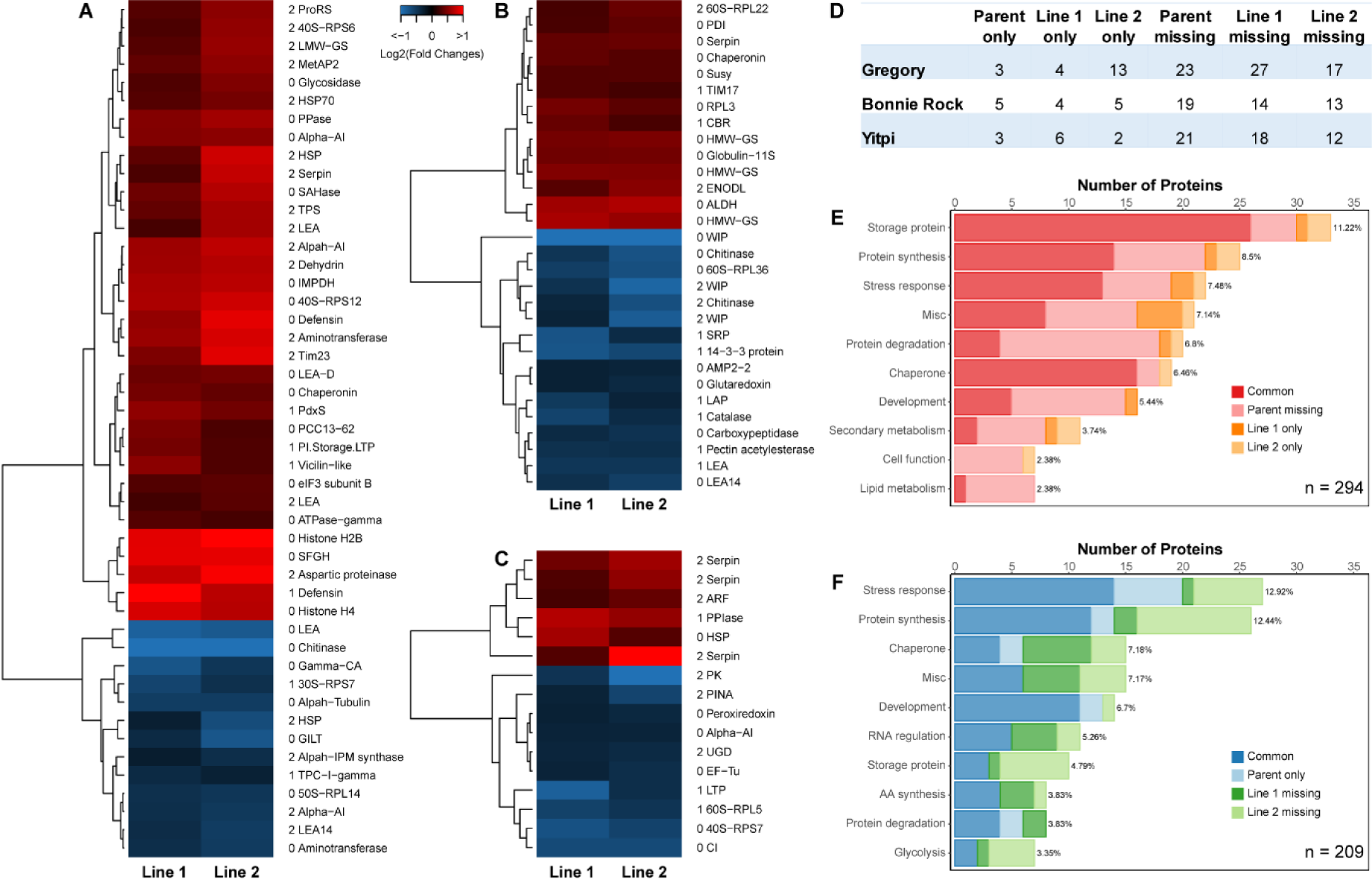
Differentially expressed proteins following integration of Ay HMW-GS across three wheat cultivars. The relative abundance for each protein was calculated through MaxQuant LFQ analysis. Fold changes in abundance for each protein were calculated by dividing the integrated line value by the parental cultivar value. Only uniquely detected proteins and commonly detected proteins that had fold change ⩾ 1.2-fold or ⩽ 0.83-fold and one-way ANOVA p-value ⩽ 0.05 were defined as differentially expressed proteins (DEPs). The full data is shown in Supplemental Table S2. Heat maps only showed DEPs that have same fold change pattern (increase or decrease) in both integrated lines. Abbreviation protein names were attached on the right side of heat maps. The first number of the protein label represents t-test p-value ⩽ 0.05 in both lines (0), in line 1 only (1) and in line 2 only (2). A, DEPs for cv. Gregory. B, DEPs for cv. Bonnie Rock. C, DEPs for cv. Yitpi. D, The number of uniquely detected proteins for three cultivars. Parent only, Line 1 only and Line 2 only represent those proteins were only detectable in parental cultivar, integrated line 1 or 2, respectively. Parent missing, Line 1 missing and Line 2 missing represent those proteins were not detectable in one of the three lines. E, Mapman functional category analysis for 294 redundant DEPs that increased in abundance were collected from all three cultivars. F, Mapman functional category analysis for 209 redundant DEPs that decreased in abundance were collected from all three cultivars. Only the top 10 functional categories are displayed. The summed proportion of all four groups for each functional category are shown.

To investigate the biological functions of this expanded set of DEPs (Fig. 3A-D), functional categories of all proteins increasing in abundance (Fig. 3E) and decreasing in abundance (Fig. 3F) were analysed. The most frequent DEPs increasing in abundance were storage proteins (33 proteins in total), which were more than 3 times the number of storage proteins decreasing in abundance (Fig. 3F). Most of the storage proteins present in higher abundance were identified from protein sets found in parent and NILs (Fig. 3A-C), while storage proteins with lower abundance were typically identified from the present/absent lists (Fig. 3D). Protein synthesis related proteins and chaperones involved in protein folding and stabilisation were similarly abundant amongst the DEPs increasing and decreasing in abundance. A higher proportion of the DEPs with low abundance were in the stress response functional category (Fig. 3F).

As the Ay introgression events in each cultivar were on the long arm of chromosome 1A, we mapped the genome location of the genes encoding DEPs to see if there were any patterns that could have arisen from this genetic location. In total, 159 DEPs from Gregory were successfully mapped, of which the number on the D sub-genome was double that on the A and B sub-genome (Supplemental Fig. S3A). Except for the apparent absence of Bonnie Rock DEPs mapped to chromosome 7B, the same pattern was also observed from the other two cultivars. The higher number of DEPs arising from genes located in the D sub-genome was consistent with the higher number of total identified proteins also mapping to the D sub-genome, and there was no significant differences between sub-genomes in terms of this proportion (Supplemental Fig. S3B). From this we concluded there was no gene location pattern of DEPs associated with the chromosome 1A integration site.

To determine which DEPs contributed most to the absolute protein abundance difference between parent and integrated lines, the 20 DEPs with the highest abundance for each parental cultivar were selected based on their absolute abundance estimated by intensity based absolute quantification (iBAQ) scores. These 20 DEPs made up 12-17% of the absolute protein abundance detected by MS in each cultivar, and consisted mainly of storage proteins, stress response proteins and ribosomal proteins (Table 2). With net increase of abundance defined via subtracting the iBAQ values of DEPs in integrated lines from its parental cultivar, the total contributions of these 20 abundant DEPs varied from 1% to 5% of total protein abundance. The serpin protein (TraesCS5D01G368900.1), the most abundant DEP in Gregory, contributed most to the total protein abundance increase in both integrated lines, being responsible for a net increase of 0.45% in line 1 and 1.59% in line 2. Likewise, the most abundant two DEPs of Bonnie Rock, alpha-amylase inhibitor (TraesCS4B01G328100.1) and 11S globulin (TraesCS1D01G067100.1), were the biggest contributors in line 1 (1.23%) and line 2 (0.98%) respectively. The heat shock protein (TraesCS3A01G033900.1) and globulin 1 (TraesCS5B01G434100.1) of cultivar Yitpi were the top contributors to integrated line 1 (0.96%) and line 2 (0.81%), respectively.

**Table 2.**
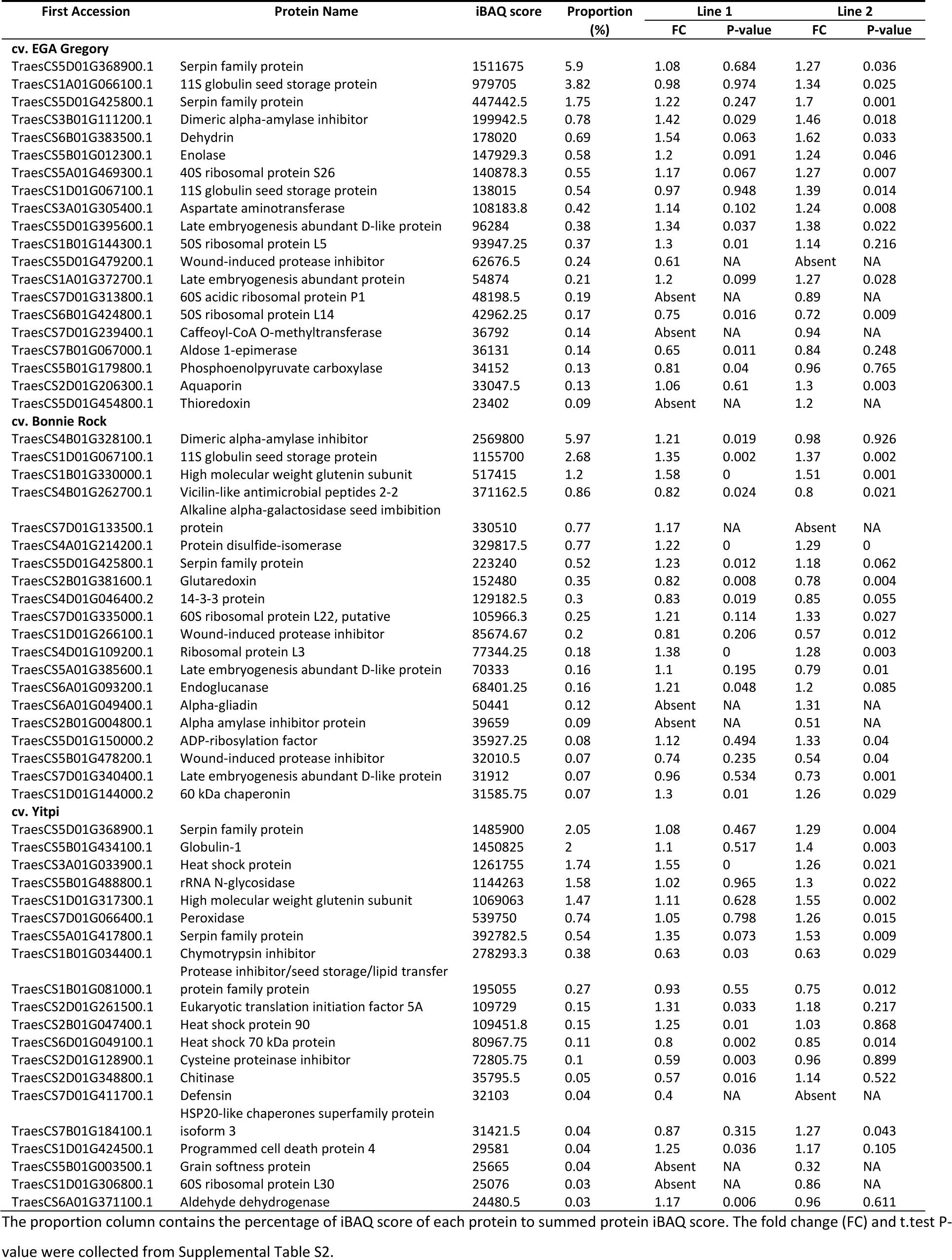
The 20 most abundant differentially expressed proteins in the Ay introgression lines of each of three wheat cultivars.

### Common changes in protein abundance induced by the introgression of 1Ay21* HMW-GS

Among the redundant set of 503 DEPs across all the cultivars shown in Fig. 3, we defined the common DEPs as those that had the same pattern of change in abundance across both integrated lines compared to their parental cultivar. In total 115 DEPs were selected including 86 that increased in abundance and 29 that decreased (Supplemental Table S3A). Nearly half of the common DEPs were storage proteins, or proteins annotated to be involved in stress response, protein synthesis, protein degradation or grain development. All the storage proteins in the set of 115 common DEPs showed higher abundance in integrated lines than in their parental cultivar (Supplemental Table S3A).

### Gene co-expression profiles of the DEPs common to introgression lines

Expression of a gene that encoded a particular protein during grain development is likely to be a significant factor in determining the final abundance of a protein product. To determine if there was co-expression of the genes encoding common DEPs, we investigated their RNA expression profiles across multiple wheat varieties, organs and growth stages. Due to the fact that wheat genes have three homologues on A, B and D subgenomes respectively, and that homologues from the D subgenome show significantly higher expression levels than those from B and A subgenomes (Ramirez-Gonzalez et al., 2018), the averaged gene expression abundance of all three homologues were used for this analysis. Transcript data for 292 homologous genes representing the common DEPs from 177 samples (23 wheat varieties, 21 organs and 37 growth stages; Wheat Expression Browser: http://www.wheat-expression.com/) were downloaded and collated and Spearman coefficient correlation analysis was performed (Supplemental Table S3). Hierarchical clustering analysis revealed two distinct gene clusters; a strong co-expression was present within one cluster while weak co-expression was found in the other (Fig. 4A). In terms of functional categories, 46 genes for storage proteins, late embryogenesis abundant proteins (LEA) and heat shock proteins (HSP) (cluster 1) were strongly co-expressed, while 68 genes encoding proteins involved in protein synthesis and degradation were not co-expressed. One exception was a carboxypeptidase that was present in cluster 1 (Fig 4A). Co-expression profiles of the genes for the 114 common DEPs were also conducted for single organs, including grain, leaf, spike and root. In comparison with the multi-organ analysis, in the grain, genes encoding storage proteins showed strong co-expression with each other but not with LEA and HSP genes, while genes for protein synthesis and degradation relevant proteins showed similar co-expression profiles (Fig. 4B). A similar gene expression profile was observed in leaf compared to grain, while less significant co-expression was found between genes of the 114 common DEPs in spike and root (Supplemental Fig. S4).

**Figure 4.**
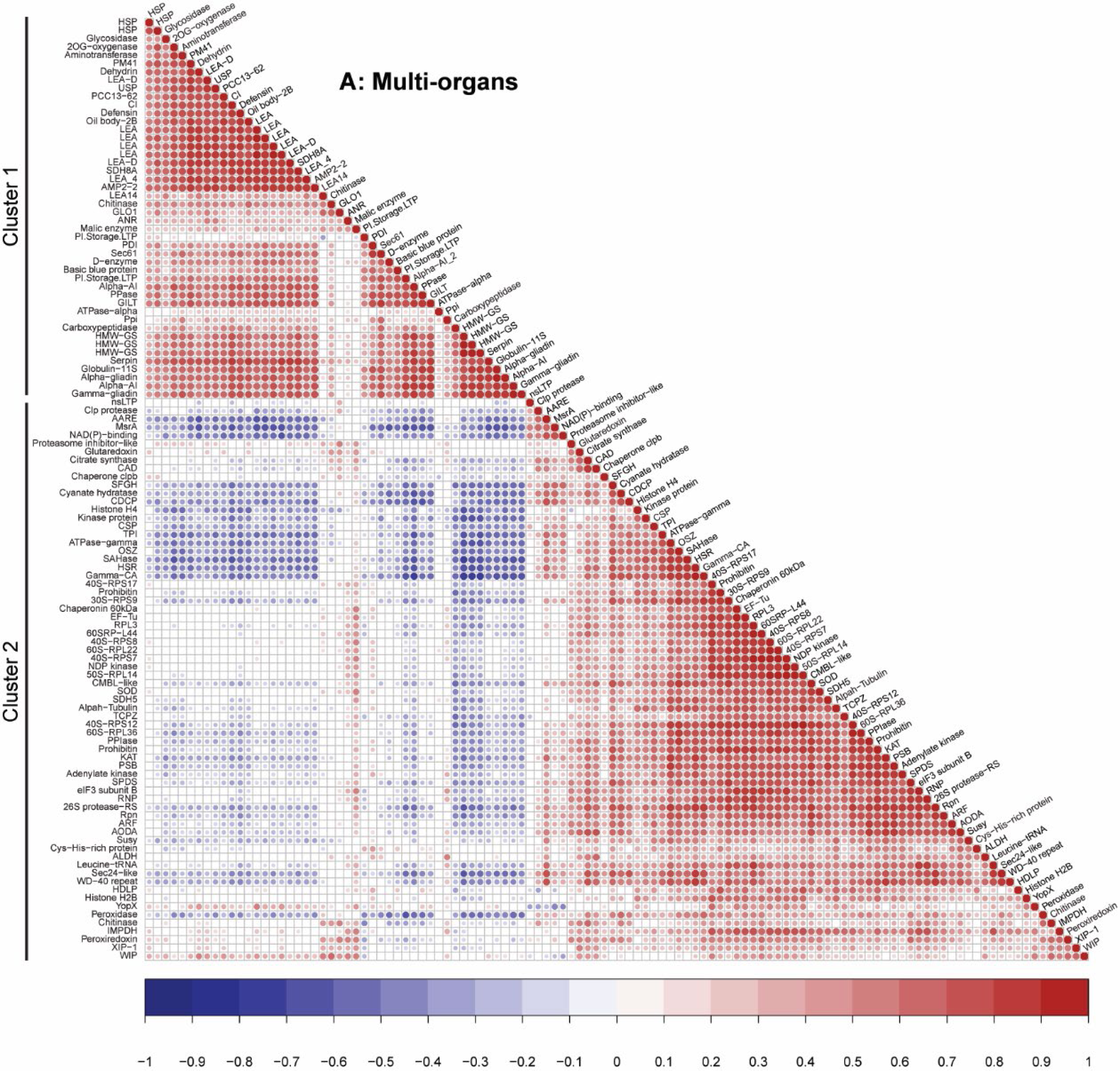
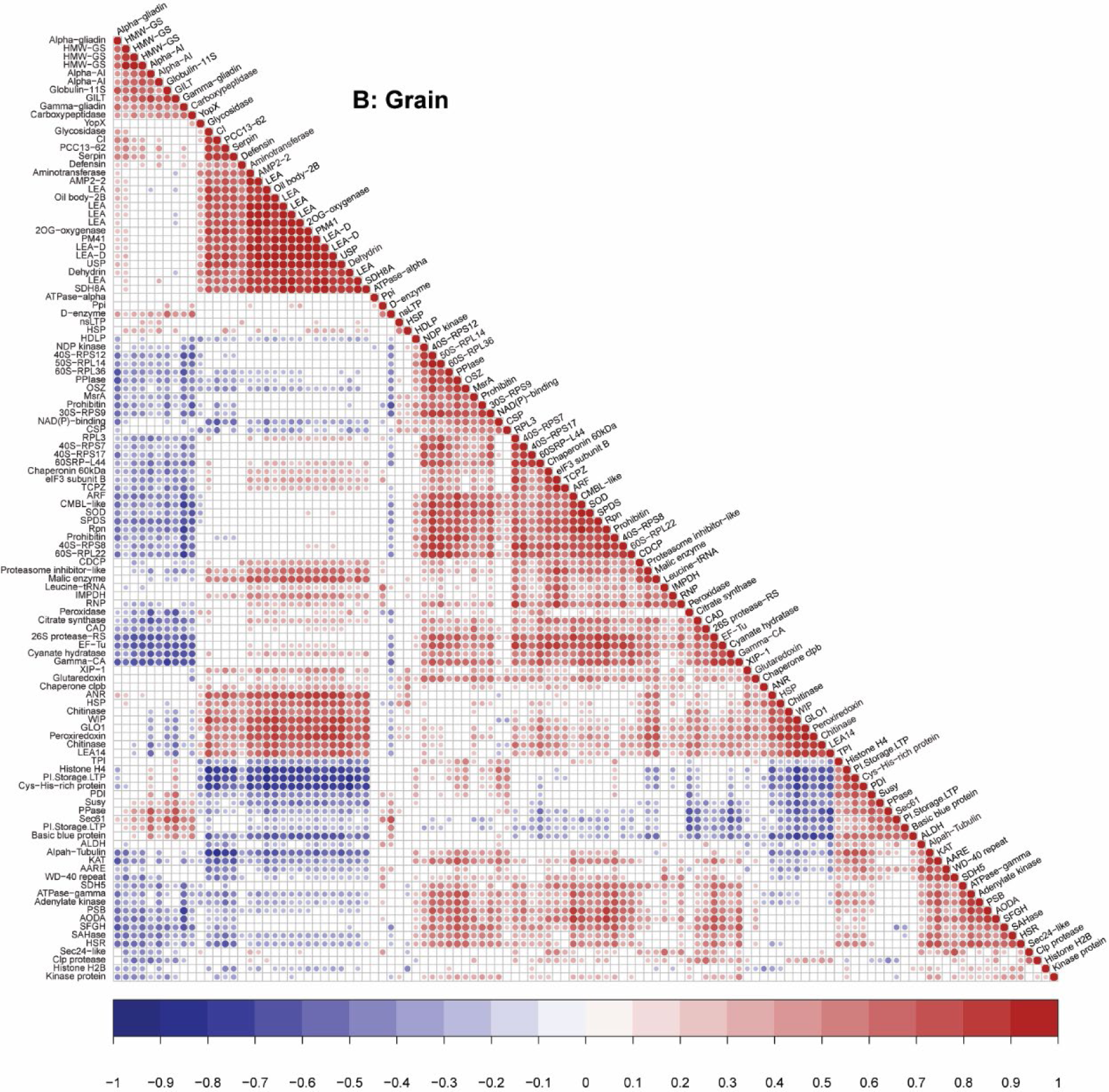
Correlation coefficients of gene expression profiles of the 114 common differentially expressed proteins. A, Coefficients of the gene co-expression profiles across 23 varieties, 21 organs and 37 growth stages (Supplemental Table S3). B, Coefficients of the gene co-expression profiles in grain only samples. The gene expression data of these general DEPs were collated from the Wheat Expression Browser (Ramirez-Gonzalez et al., 2018). Spearman correlation analysis and hierarchical clustering for both columns and rows was used in data visualization. The abbreviation protein names are shown. Pairs with P-value ⩾ 0.05 were removed leaving blank cells on the plot.

### Analysis of *cis*-acting elements present in associated wheat gene promoter regions

As gene co-expression can indicate co-transcriptional activation by common transcription factors (TF), common TF binding sites (TFBSs) may be present in upstream promoter regions of the genes of interest in this analysis. To assess this, the *cis*-acting regulatory elements located in the upstream promoter region of gene homologues representing common DEPs and non-DEPs were analysed using plantCARE (Supplemental Table S4A-C). In total, 140 *cis*-elements belonging to 7 sub-categories were identified. Most of these *cis*-elements are associated with physiological processes including light response, developmental regulation, hormonal response and environmental stress response. A total of 894 genes that encode non-DEPs identified in grain proteomes were used as control. Statistical analysis suggested that promoter related elements (TATA-box and CAAT-box), and the majority of light responsive (G-box, G-Box, Box-4, GATT-motif, GT1-motif and Pc-CMA2c) and hormone responsive (ABRE, P-box and AuxRR-core) *cis*-elements were significantly enriched in the DEP set compared to the control non-DEP set (Table 3). *Cis*-elements related to development (circadian and MBS) and environmental stress response (GC-motif and Sp1) showed the opposite pattern with higher enrichment in the non-DEP background genes than in common DEP genes. This analysis was repeated using a randomly sampled set of 1000 wheat genes as a background dataset, and that returned a result highly consistent with using the 894 non-DEP gene set as control (Supplemental Table S4E). Comparison of the two background datasets showing no statistical differences in *cis*-element enrichment between them (Supplemental Table S4F).

**Table 3.** *Cis*-acting regulatory element analysis of upstream 1500bp promoter region of genes encoding differentially expressed proteins in Ay introgression lines.

| Element Name | No. Cluster 1 |  | No. Cluster 2 |  | No. Background |  | P-value (Fisher's Exact Test) |  |  | Function category |
| --- | --- | --- | --- | --- | --- | --- | --- | --- | --- | --- |
|  | Total | Mean | Total | Mean | Total | Mean | Cluster 1 vs Background 1 | Cluster 2 vs Background 1 | Cluster 1 vs Cluster 2 |  |
| Cluster 1 enriched *cis*-element sites |  |  |  |  |  |  |  |  |  |  |
| TATA-box | 1504 | 15.67 | 2274 | 15.79 | 12265 | 13.72 | 0 | 0 | 0.129 | Promoter related elements |
| WUN-motif | 35 | 0.36 | 30 | 0.21 | 186 | 0.21 | 0.006 | 0.921 | 0.043 | Environmental stress-related elements |
| G-box | 308 | 3.21 | 336 | 2.33 | 2410 | 2.7 | 0.013 | 0.024 | 0 | light responsive elements |
| G-Box | 117 | 1.22 | 121 | 0.84 | 852 | 0.95 | 0.027 | 0.281 | 0.012 | light responsive elements |
| Box 4 | 73 | 0.76 | 69 | 0.48 | 457 | 0.51 | 0.004 | 0.751 | 0.013 | light responsive elements |
| GATT-motif | 5 | 0.05 | 1 | 0.01 | 9 | 0.01 | 0.009 | 1 | 0.045 | light responsive elements |
| ABRE | 454 | 4.73 | 400 | 2.78 | 3109 | 3.48 | 0 | 0 | 0 | Hormone responsive elements |
| P-box | 37 | 0.39 | 37 | 0.26 | 225 | 0.25 | 0.029 | 0.857 | 0.124 | Hormone responsive elements |
| Cluster 2 enriched *cis*-element sites |  |  |  |  |  |  |  |  |  |  |
| CAAT-box | 1823 | 18.99 | 2888 | 20.06 | 16178 | 18.1 | 0.263 | 0 | 0 | Promoter related elements |
| AT-rich element | 7 | 0.07 | 16 | 0.11 | 55 | 0.06 | 0.67 | 0.037 | 0.397 | Site-binding related elements |
| GT1-motif | 45 | 0.47 | 91 | 0.63 | 445 | 0.5 | 0.649 | 0.032 | 0.066 | light responsive elements |
| Pc-CMA2c | 1 | 0.01 | 5 | 0.03 | 6 | 0.01 | 0.518 | 0.011 | 0.411 | light responsive elements |
| AuxRR-core | 10 | 0.1 | 27 | 0.19 | 104 | 0.12 | 0.875 | 0.029 | 0.095 | Hormone responsive elements |
| Non-DEPs enriched *cis*-element sites |  |  |  |  |  |  |  |  |  |  |
| motif I | 9 | 0.09 | 1 | 0.01 | 46 | 0.05 | 0.113 | 0.017 | 0.002 | Development related elements |
| circadian | 6 | 0.06 | 4 | 0.03 | 111 | 0.12 | 0.088 | 0.001 | 0.335 | Development related elements |
| MBS | 72 | 0.75 | 68 | 0.47 | 579 | 0.65 | 0.323 | 0.018 | 0.012 | Environmental stress-related elements |
| GC-motif | 58 | 0.6 | 75 | 0.52 | 673 | 0.75 | 0.081 | 0.003 | 0.537 | Environmental stress-related elements |
| Sp1 | 70 | 0.73 | 105 | 0.73 | 1107 | 1.24 | 0 | 0 | 0.817 | light responsive elements |
| 3-AF1 binding site | 0 | 0 | 4 | 0.03 | 34 | 0.04 | 0.045 | 0.813 | 0.15 | light responsive elements |
| A-box | 63 | 0.66 | 113 | 0.78 | 875 | 0.98 | 0.001 | 0.041 | 0.167 | Hormone responsive elements |

## DISCUSSION

Introgression of Ay genes has been trialled for grain protein content improvement in wheat breeding programmes worldwide (Payne, 1983; Waines and Payne, 1987; Rogers et al., 1997; Shewry and Halford, 2002). Many studies have been conducted to either characterize Ay gene structures or explore its economic potential in modern wheat breeding programmes (Margiotta et al., 1996). These reports have mainly focused on gene discovery and gene structure characterization (Bai et al., 2004; Jiang et al., 2009; Gutierrez et al., 2011; Bi et al., 2014; Yu et al., 2019), and the influence of Ay HMW-GS introgression on agronomic and quality traits, including grain yield, grain protein content, storage protein composition and dough quality (Rogers et al., 1997; Roy et al., 2018; Roy et al., 2020). By studying the proteome of Ay integrated near isogenic lines we have provided insights into the nature of the changes in the wider grain protein profile of bread wheat induced by Ay introgression events and propose a model to explain the molecular events observed.

### Introgression of the expressed *Ay* gene changes more than the abundance of *Ay* itself

The initial intent of cross breeding the *Glu-1Ay* gene into hexaploid wheat was to improve the grain protein content and quality by addition of an extra Ay HMW glutenin subunit. Previous work has proved that the Ay HMW-GS integrated lines have higher grain protein content (Rogers et al., 1997; Roy et al., 2018). However, it was unknown whether the increased protein abundance was solely from the Ay HMW-GS protein itself, or if there were other wheat grain proteins contributing to the protein content improvement. Our results showed that the integrated lines had significantly higher protein content compared to parental cultivars as well as an increase of gluten content (Table 1). Calculations based on the contribution of HMW-GS to total gluten (Zorb et al., 2018) showed that the total increase in all gluten proteins represented on average only 52% of the total protein content increase. The proteome data showed that approximately a quarter of the proteins identified in wheat grains by MS significantly changed in their abundance in Ay integrated lines (Fig. 3). The most abundant 20 DEPs of each cultivar (predominantly storage proteins, stress response proteins and ribosomal proteins) contributed most of the remaining 48% increase of total protein abundance (Table 2). The MapCave functional category analysis showed these common DEPs belonged to 36 different functional categories, indicating it is not a single system or a discrete part of the cellular machinery that is altered following Ay introgression. The genomic locus analysis showed that DEPs arise from genes that are relatively evenly distributed across the 21 chromosomes of bread wheat, so changes in gene expression that might be driving these changes in protein abundance are not tightly associated with the introgression event on chromosome 1A. It appears expression of Ay HMW-GS has a significant downstream impact resulting in alterations to the wheat grain proteome which causes the significant increase in grain protein. The *cis*-element analysis conducted provides support for Ay HMW-GS integration-dependent activation of genes encoding common DEPs because their promoters were enriched with TATA-box and light response elements (Table 3). Specific downstream targets involved in ATP synthesis, amino acid synthesis and degradation, protein folding and RNA regulation (Supplemental Table S2H and I) can be treated as valuable protein candidates for future breeding programmes of grain protein content and bread-making quality improvement. However, it was the changes in abundance of three major groups of proteins that dominated the protein response; storage proteins, proteins of the protein synthesis machinery, and proteins involved in protein stability and degradation systems.

### Ay HMW-GS expression triggers changes in the abundance of other storage proteins

A total of 43 storage proteins, other than Ay HMW-GS, were defined as DEPs in this analysis, among which 33 had higher abundance and 10 had lower abundance in Ay integrated lines. All the HMW-GS (7 proteins), Globulin-11S (5 proteins) and all but one of the Serpins (10 proteins) increased in abundance. These HMW glutenin subunits are key determinants in dough quality formation and bread-making quality and may be important contributors to the quality traits attributed to Ay introgression lines (Shewry and Halford, 2002; Roy et al., 2020). Globulin-11S is the main form of the water-soluble globulin proteins that account for approximately 10% of grain storage protein (Hellemans et al., 2018; Zorb et al., 2018). Although the role of globulin-11S in bread-making quality is not very clear, evidence indicates that it is able to form inter- and intra-chain disulfide links and partially contributes to variations found in dough quality (Inquello et al., 1993; Jung et al., 1997; Hellemans et al., 2018). Serpins make up 4% of total grain protein and may contribute to the increase of grain protein content through their role in inhibiting proteases, thereby increasing the stability of storage proteins (Ostergaard et al., 2000; Cane et al., 2008). Serpins can influence bread-making quality either via modifying the molecular structure of prolamin storage proteins or by forming intermolecular disulfide bridges between serpins, and between serpins and β–amylase proteins (Roberts and Hejgaard, 2008). Taken together, the accumulation of these three protein families (especially HMW-GSs) are likely to be the major driving force in the improvement of grain protein content and bread-making quality observed in Ay integrated lines. Better bread-making quality in Ay integrated lines in the Bonnie Rock genetic background have been reported (Roy et al., 2018), and it was in this background that the largest and most consistent increases in gluten content and HMW-GSs were observed in this study (Fig 1 and 3, and Table 1).

### Ay HMW-GS induced selective changes in protein synthesis machinery

As the basic functional units of protein synthesis machinery, ribosomes and their function throughout grain development will be important to higher grain protein content. Eukaryotic ribosomes are composed of small 40S and large 60S subunits containing up to 100 different proteins. Ribosomes are initially assembled in the nucleolus and then released into the cytoplasm as pre-ribosomal particles (Fromont-Racine et al., 2003; Panse and Johnson, 2010). In plants, the specific isoforms of a given ribosome protein can differ between tissue types (Salih et al., 2020) and are exchanged during environmental stress like cold and oxidative stress (Bailey-Serres and Freeling, 1990; Byrne, 2009; Salih et al., 2020). Ribosome composition is critical for the speed and efficiency of protein synthesis, especially of proteins containing rare codons (Novoa and Ribas de Pouplana, 2012). In this study, 43 specific ribosomal proteins showed significant changes in abundance, even though the overall net increase in amount was minor within Ay integrated lines compared to parental cultivars (Supplemental Table S2H and I). Non-ribosomal proteins that support translation, such as the eukaryotic translation initiation factor 3B and 5A and elongation factor 1 alpha were DEPs that increased in abundance (Supplemental Table S2H and I). These complicated patterns might imply that protein synthesis machinery regulation is initiated in Ay lines that may support changes in storage protein synthesis.

### Ay HMW-GS induced changes to the protein folding, stability and proteolysis systems in grains

There is no doubt that keeping newly assembled proteins stable can be as important as making new proteins to achieve higher protein content (Fink, 1999; Lee et al., 2003). Summing up the total abundance increases and decreases for 34 molecular chaperones showed a net accumulation of 2.3-fold. During the grain maturation period, protein aggregation is triggered by desiccation, hence having more molecular chaperone proteins will better prevent irreversible protein aggregation and help wheat grain withstand severe desiccation (Liberek et al., 2008). The accumulation of late embryogenesis abundant (LEA) proteins usually occurs during mid to late embryogenesis and correlates with the acquisition of seed desiccation tolerance (Wise and Tunnacliffe, 2004). Summing up the net impact of 30 changes to LEA protein abundance (16 increases and 14 decreases) showed a net increase of 1.7-fold. The combination of accumulation of chaperones and LEA in Ay integrated lines may benefit higher grain protein content through improving the stability of storage proteins during grain development. To maintain protein homeostasis, proteolysis is also required to remove damaged, mis-folded and dysfunctional proteins. The ubiquitin-proteasome system is the principal mechanism responsible for degrading short-lived regulatory proteins and soluble mis-folded proteins (Zhang et al., 2007; Marshall and Vierstra, 2019). Autophagy, by contrast, can eliminate larger protein complexes and insoluble protein aggregates and it has been reported to be involved in the degradation of storage proteins during seed germination (Vanderwilden et al., 1980; Toyooka et al., 2001; Bassham, 2007). Our results indicated that proteasomal proteins, such as the 26S protease regulatory subunit and proteasome β subunit, showed an increase pattern in abundance while most lysosomal proteases like carboxypeptidase and leucine aminopeptidase decreased in abundance in Ay introgression lines (Supplemental Table S2H and I). Thus, we could hypothesize that the autophagy-lysosome system was suppressed to stabilize newly synthesized storage proteins during the accumulation of storage protein in Ay lines. In contrast, increased 26S proteasome system could recycle nutrients from regulatory or incorrectly folded proteins to optimise protein abundance.

### A putative model of *Glu-Ay* induction of high grain content and bread-making quality

Combining our results and previous reports, we proposed a model that can now be tested of how introgression of the 1Ay21* HMW-GS improves wheat grain protein content and bread-making quality (Fig. 5). This proposal begins with the expression of the extra Ay gene taking the total number of HMW glutenin subunits from 5 to 6. Increased HMW-GS content is deposited into protein storage vacuoles through either the Golgi apparatus into the vacuole or the protein body accumulating directly within the lumen of the ER (Muntz, 1998; Tosi et al., 2009). By unknown mechanisms, this increased rate of protein body accumulation activates transcription factors that bind sites in the promoter regions of a suite of genes that already form a developmental gene expression program to enhance storage proteins and protein synthesis regulatory proteins in wheat grains. Ribosomal composition changes may then fulfil the higher demand for protein synthesis and more molecular chaperones assist nascent proteins to fold properly and be stored. Due to the protection from higher abundances of chaperones and LEA, other storage proteins are deposited into protein storage vacuoles with enhanced stability, which led to less need of the lysosome system and a broad increase in the abundance of HMW-GS, Globulin-11S and Serpins. Boosting of the ubiquitin-proteasome system recycles nutrients from unwanted proteins, providing a positive reinforcement and resources for more storage protein synthesis and deposition. As a consequence of these concerted actions, the grain protein content for Ay HMW-GS introgressed lines increases, leading to improvement of dough quality and bread-making quality because of the alteration of gluten composition.

**Figure 5.**
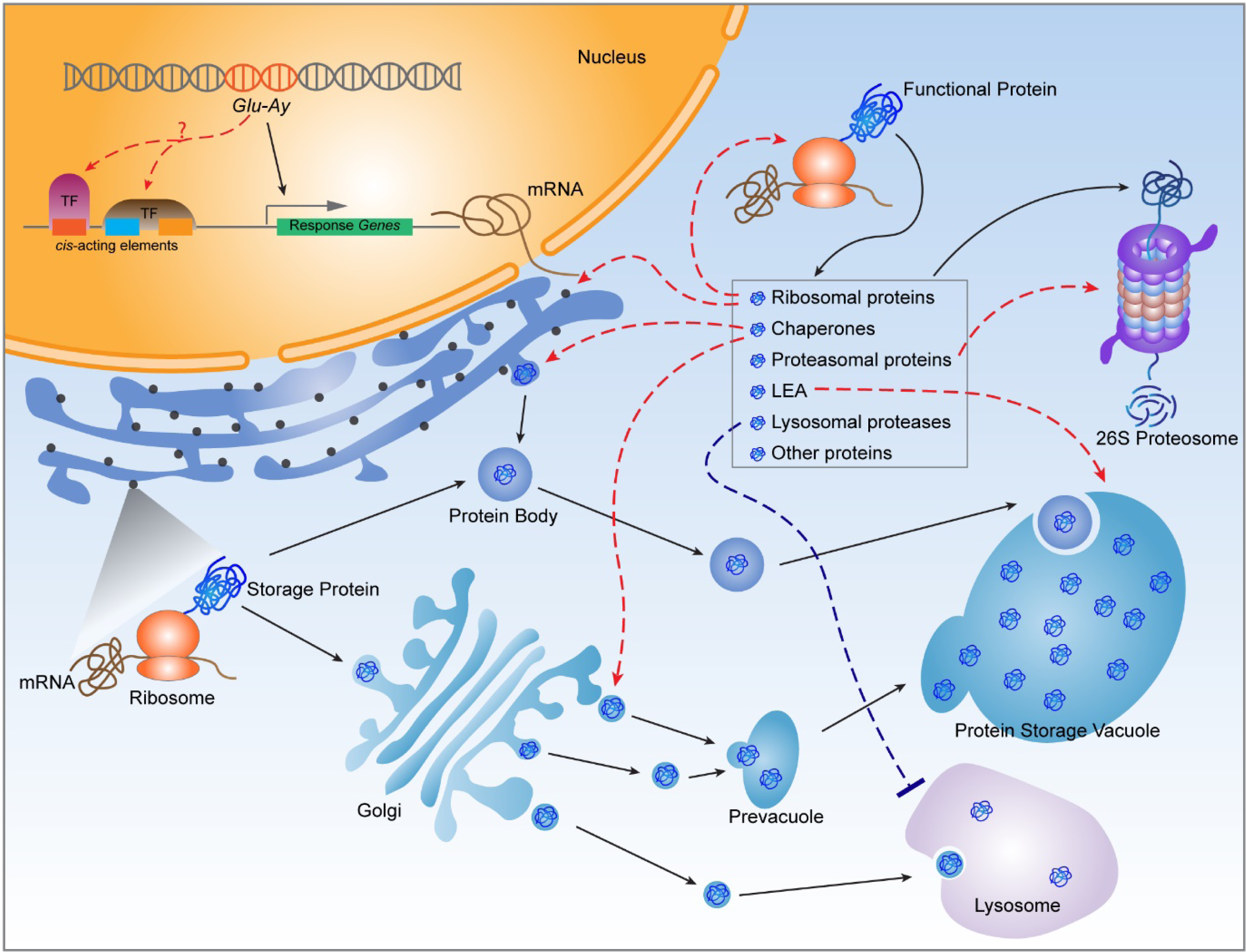
Model of a putative pathway for grain protein accumulation triggered by the introgression of 1Ay 21* HMW-GS. Black arrows represent general protein trafficking pathway; red dashed arrows represent the positive regulation effects while the blue dashed flat-headed arrow represent negative regulation effects. TF: Transcription factor; ER: Endoplasmic reticulum.

## MATERIALS AND METHODS

### Plant materials

The introgression of the expressed *Glu-1Ay* gene from an Italian wheat line N11 into Australian wheat cultivars EGA Gregory, EGA Bonnie Rock and Yipti, were completed using conventional cross breeding processes as previously reported (Roy et al., 2018; Roy et al., 2020). Near isogenic lines (NILs) for each cultivar were developed through the above cross breeding process and materials including one parental cultivar and two integrated sister lines were selected for this study. The NILs and their corresponding parental cultivars were grown at the experimental field station of Department of Primary Industries and Regional Development (DPIRD) at South Perth, Western Australia in 2017 following a Complete Randomised Design (CRD) with three replicates. Grains were harvested from the field plot at full maturity, and 100g of subsamples of each replicate was prepared for phenotyping. Grain protein content (GPC) and gluten were measured using Near Infrared Transmission Spectrophotometry through the “CropScan 3000F Flour and Grain Analyser” (Next Instruments, Condell Park, NSW-2200, Australia). Each measurement was taken by averaging the data of 10 individual scanning events.

### Grain protein sample preparation

Protein extraction involved a very fine grinding of whole grains with excess reductant and a high SDS concentration in order to combine the so called extractable polymeric protein (EPP) and unextractable polymeric protein (UPP) fractions from wheat grains (Vensel et al., 2014) into a single sample. Samples of 4 biological replicates for each line and 20 seeds per replicate were ground with mortar and pestle under liquid nitrogen. 200 mg fine ground power were weighed for further total protein extraction using a chloroform/methanol precipitation approach (Wessel and Flugge, 1984). Briefly, samples were thoroughly mixed with 400 µL extraction buffer that contains 125 mM Tris–HCl pH 7.5, 7% (w/v) sodium dodecyl sulfate, 10% (v/v) b-mercaptoethanol, 0.5% (w/v) PVP40 and Roche protease inhibitor cocktail (1 tablet per 50 ml of extraction buffer, Roche). After rocking on ice for 10 min, mixture was centrifuged at 10,000 x g, 4°C for 5 min to separate the supernatant and pellet. About 200 µL supernatant was then moved into a new tube following by a precipitation step via adding 800 µL methanol, 200 µL chloroform and 500 µL distilled deionized water. With another centrifugation at 10,000 x g, 4°C for 5 min, the upper aqueous phase was carefully removed and the protein pellet was washed twice with 500 µL methanol. The protein pellet was then incubated with 1 mL 90% (v/v) acetone at -20°C for at least one hour for two times. After drying at room temperature, the dried protein pellet was resuspended with 40-100 µL resuspension buffer (50 mM Ammonium bicarbonate, 1% sodium dodecyl sulfate and 10 mM DL-dithiothreitol). Amido black quantification method was applied for protein concentration measurement (Schaffner and Weissmann, 1973).

For protein digestion, 200 µg of proteins first incubated with 20 mM DL-dithiothreitol for 30 min in the dark, which then followed by a second incubation with 20 mM iodoacetamide for 30 min in the dark. After diluting the concentration of sodium dodecyl sulfate to 0.1% (v/v), the overnight protein digestion was conducted using trypsin (sigma, USA) with the trypsin-protein ratio at 1:50. Sodium dodecyl sulfate removal was performed using the method reported by Yang et al with minor modifications (Yang et al., 2012). Briefly, peptide samples were injected into an off-line Agilent 1200 series HPLC configured with two J4SDS-2 guard columns (PolyLC, Columbia, USA). Peptides were eluted by 60% (v/v) acetonitrile at the time window from 2-5 min, and target peptides were collected from 3.7 min to 4.5 min. After drying down in a vacuum centrifuge, dried peptides were resuspend with 5% (v/v) acetonitrile and 0.1% (v/v) formic acid in water following a further filtering with 0.22 µm centrifugal filter units (Millipore, USA) to remove any undissolved pellets. Purified peptide suspensions were stored at -80°C for further use.

### Mass spectrum acquisition

Purified peptide suspension was injected into a HPLC-chip (Polaris-HR-Chip-3C18) using a capillary pump with a flow at 1.5 μL/min, 2 μL. Peptides were eluted from the C18 column and into an online Agilent 6550 Q-TOF (Agilent Technologies, USA). A two-hour gradient generating by a 1200 series nano pump (Agilent Technologies, USA) with the nano flow at 300 nl/min was conducted for LC-MS/MS. The elution gradient started with 5% (v/v) solution B (0.1% (v/v) formic acid in acetonitrile), following gradients 5 to 6% in 6 min, 6-22% in 84 min, 22-35% in 5 min, 35-90% in 3 min, remained 90% for 4 min, 90-5% in 2 min. Parameters setting for mass spectrum acquisition were previously reported by Duncan et al., (Duncan et al., 2017). Briefly, data dependent mode and a scan range from 300 to 1750 mz was used for MS acquisition. MS data was collected at eight spectra per second and MS/MS data was collected at four spectra per second. Ions were dynamically excluded for 6 sec following fragmentation. In total, MS data for 36 samples were successfully collected. The primary MS data files are available via ProteomeXchange with identifier PXD021706.

### Label free quantification of precursor ion intensities

To obtain protein relative and absolute abundances, label free quantification was performed using MaxQuant (version 1.6.1.0) (Cox and Mann, 2008). For protein relative abundance, 36 Agilent .d files collected from mass spectrometry analysis for 9 lines and 4 bio-replicated per line were upload to MaxQuant and searched against the wheat protein database (IWGSC, http://www.wheatgenome.org/, version 1.0, 137029 sequences) with reversed decoy sequences automatically attached by MaxQuant (Appels et al., 2018). The search was conducted under general LFQ model with 20 ppm for mass tolerance and 1% FDR. The LFQ-intensity of each biological sample was used as relative abundance for downstream data analysis. Protein groups were considered if they were identified ≥2 replicates with at least 6 independently quantified peptides from the 4 bio-replicates. In order to further improve data quality, an additional filter of relative standard deviation by mean was also used, in which only those protein groups having rsd ≤ 30% or sd ≤ 30% of overall mean abundance were retained, and ∼15% of data from each genotype was trimmed off (Supplemental Fig. S1). In terms of the estimation of protein absolute abundance, MaxQuant search with the same parameter sets as mentioned above and the additional iBAQ (intensity Based Absolute Quantification) model was performed.

To detect the presence of 1Ay21* HWM-GS in integrated lines and to estimate its absolute abundance along with the other five HMW-GSs, the iBAQ search of each cultivar against a dedicated 6 HMW-GSs sequences database collected from UniProt (https://www.uniprot.org/) was conducted. The detailed HMW-GSs composition for each cultivar was summarized in Supplemental Table S1A. The fold change in abundance was calculated using the iBAQ protein abundance of integrated lines divided by its corresponding parental cultivar, while one-way ANOVA with post-hoc Tukey HSD test was performed for statistical analysis. Proteins with fold change ≥ 1.2 or ≤ 0.83 and p-value ≤ 0.05 were defined as differentially expressed proteins (DEPs). Uniquely present or absent proteins were also considered as DEPs. Common DEPs were defined as DEPs with the same fold change direction in both integrated lines compared to a parental cultivar. 115 common DEPs were identified including 48, 38, and 29 DEPs for Gregory, Bonnie Rock, and Yitpi, respectively (Supplemental Table S3A).

### Gene co-expression analysis

Gene expression data under non-stress conditions were collated from the Wheat Expression Browser website (http://www.wheat-expression.com/) and used for gene co-expression profiles of the common DEPs (Ramirez-Gonzalez et al., 2018). The first protein IDs of the protein groups, as the representative protein ID of the group, were used for transcript data collection. The averaged gene expression abundance of A, B and D genome homologues were calculated and used in this analysis. The gene co-expression relationship was estimated by Spearman coefficient correlation, and conducted on multi-organ, grain only, leaf only, spike only and root only datasets. Two of the 115 genes were homologues resulting to only 114 triads remaining for gene co-expression analysis.

### Identification of *cis*-acting regulatory elements on promoter regions

The 1500-bp upstream promoter sequences starting from the translation initiation codon were downloaded from Ensembl Plants (https://plants.ensembl.org/index.html). In total 107 of 144 common DEPs encoding genes were successfully collected. The 1500-bp upstream promoter sequences for two background data sets were also collected. Background dataset 1 included 894 genes representing 1827 redundant non-DEPs over three cultivars, while the background dataset 2 contained 964 genes that were randomly sampled from the IWGSC database (137052 total genes, sampled 1000 genes and 36 genes failed promoter sequence collection). These promoter sequences were then submitted to plantCARE (http://bioinformatics.psb.ugent.be/webtools/plantcare/html/) for *cis*-element identification (Lescot et al., 2002).

### Data statistical analysis and visualization

Data processing and statistical analysis was performed using R (version 3.5.1), including PCA, one-way ANOVA, Fisher’s Exact Test and coefficient correlation.

## ACKNOWLEDGEMENT

H.C was supported by Research Training Program Fee Offset-International Student and UWA Safety-Net Top-Up Scholarship. This work was support by Australian Research Council funding to AHM (CE140100008) and Grain Research and Development Corporation funding to WM (UMU00036).

## SUPPORTING DOCUMENTS

**Supplemental Figure S1**. Peptide mass spectral data quality controls.

**Supplemental Figure S2**. Volcano plot for commonly detected proteins.

**Supplemental Figure S3**. The distribution of DEPs on wheat chromosomes.

**Supplemental Figure S4**. The gene co-expression profiles of 114 common DEPs on individual organ.

**Supplemental Table S1**. The list of absolute abundance (iBAQ) of 6 HMW-GSs for each of the three cultivars.

**Supplemental Table S2**. The list of relative abundance of all quantified proteins for each of the three cultivars.

**Supplemental Table S3**. The gene co-expression data of the homologous genes of the 114 common DEPs.

**Supplemental Table S4**. The cis-acting regulatory element data of the homologous genes of the 107 common DEPs.

